# Thyroid hormone promotes fetal neurogenesis

**DOI:** 10.1101/2025.05.14.654075

**Authors:** Federico Salas-Lucia, Sergio Escamilla, Amanda Charest, Hanzi Jiang, Randy Stout, Antonio C. Bianco

**Author notes:** Corresponding author: Federico Salas-Lucia, PhD, Section of Adult and Pediatric Endocrinology and Metabolism, Department of Medicine, University of Chicago Medical Center, 5841 S. Maryland Ave. MC1027, Room M267 | Chicago, IL 60637.

## Abstract

Maternal low thyroxine (T4) serum levels during the first trimester of pregnancy correlate with cerebral cortex volume and mental development of the progeny, but why neural cells during early fetal brain development are vulnerable to maternal T4 levels remains unknown. In this study, using iPSCs obtained from a boy with a loss-of-function mutation in MCT8—a transporter previously identified as critical for thyroid hormone uptake and action in neural cells—we demonstrate that thyroid hormones induce transcriptional changes that promote the progression of human neural precursor cells along the dorsal projection trajectory. Consistent with these findings, single-cell, spatial, and bulk transcriptomics from MCT8-deficient cerebral organoids and cultures of human neural precursor cells underscore the necessity for optimal thyroid hormone levels for these cells to differentiate into neurons. The controlled intracellular activation of T4 signaling occurs through the transient expression of the enzyme type 2 deiodinase, which converts T4 into its active form, T3, alongside the coordinated expression of thyroid hormone nuclear receptors. The intracellular activation of T4 in NPCs results in transcriptional changes important for their division mode and cell cycle progression. Thus, T4 is essential for fetal neurogenesis, highlighting the importance of adequate treatment for mothers with hypothyroidism.

## INTRODUCTION

Thyroid hormones are important for a normal pregnancy, both for the mother and the developing fetus. While it is generally known that thyroid hormones are critical for brain development (1), much attention has been focused recently on the first trimester of the pregnancy, a period in which the fetal thyroid is not yet functional and the fetal supply of thyroid hormones depends on the maternal-fetal transport across the placenta (2, 3). Indeed, maternal T4 can be found in the human fetus before their thyroid gland becomes functional (4). Elegant studies that utilized radioactive tracers demonstrated that ^125^I-T4 but not ^125^I-T3 injected in pregnant dams can reach the fetal brain (5). These findings elevate the role played by maternal T4 levels during early pregnancy. Unfortunately, low maternal T4 levels during the first trimester are not uncommon, occurring in ∼25% of all pregnancies (6). Later in life, children from these pregnancies may exhibit reduced cerebral cortex volume and lower IQ (7), suggesting that low maternal T4 levels can be detrimental for early brain development.

T4 is a prohormone with little biological activity, but it can be activated by conversion to T3 (8). Therefore, the dependence on T4 and the finding of ^125^I-T3 in the fetal brain after mothers were injected with ^125^I-T4 implies that fetal deiodinases can activate T4 to T3 and trigger thyroid hormone action. The type 2 deiodinase (DIO2) is a key enzyme producing T3 in the fetal brain (9), and is expressed at high levels in the cerebral cortex during early to mid-gestation. This explains why, at week 14, T3 levels in the fetal cerebral cortex are higher than those in adult brain despite considerably lower circulating T3 levels in the fetus (10). Therefore, early DIO2 expression in the fetal brain is likely to ensure that T3 signaling is initiated and sustained during early brain development.

In fact, as early as gestational week 16, *DIO2* is expressed in human (and mice) neural precursor cells (NPCs) (11, 12)−the source of most cortical neurons−supporting a role for DIO2 in early neurogenesis (the process by which new neurons are formed in the brain). In agreement, our previous studies using iPSC-derived cortical organoids (COs) to model the first trimester of human fetal brain development (6.5 to 14 gestational weeks) demonstrated that a reduction in thyroid hormones transport into fetal neural cells results in abnormal neurogenesis (13). These abnormalities were traced back to alterations in the division mode of NPCs, which determines whether NPCs keep proliferating or start their differentiation into neurons. The presence of DIO2 and the relevance of thyroid hormones for the division mode of NPCs is very intriguing and begs the question: Can T4 (via the DIO2 pathway) act in NPCs to trigger developmental programs in these cells, ultimately regulating their proliferation and neuronal differentiation?

Here, we leveraged single-cell, spatial, and bulk transcriptomics to interrogate how T4 affects the molecular and cellular characteristics of the NPCs and their potential to differentiate into neurons. First, we modeled suboptimal thyroid hormone levels during the early stages of cerebral cortex development using COs generated from iPSC prepared with cells from a boy with a loss of function mutation in MCT8. MCT8 is a TH transporter we previously identified as critical for TH uptake and action in neural cells (13), and COs with a non-functional MCT8 exhibited an arrested progression of the NPCs into the dorsal projection trajectory (i.e., NPCs → intermediate progenitors → projection neurons (14)). Then, we used enzymatic assays to demonstrate that NPCs are equipped with MCT8 and DIO2 and that these components work together to rapidly trigger thyroid hormone signaling by taking up T4 and building up intracellularly generated T3 in the NPCs. The D2-generated T3 triggers genetic programs in NPCs related to their cell cycle progression and is critical for neuronal differentiation. These results reveal a previously unappreciated role of T4 in promoting early fetal cerebral cortex development, providing a mechanistic explanation as to why maternal T4 is so important for fetal cerebral cortex development during the first trimester of the pregnancy.

## RESULTS

### The progression of the dorsal projection trajectory of the human cerebral cortex depends on thyroid hormones

To understand the mechanistic basis of how maternal hypothyroxinemia (low T4 levels) affects early cerebral cortex development (7), we prepared 50-day-old (D50) COs using MCT8-deficient iPSC prepared with cells from a 6-year-old boy with the Allan-Herndon-Dudley Syndrome (carrying the missense mutation P321L). iPSC prepared with cells from the child’s father were used as controls. All COs were prepared in medium containing 20 pM free T3. A pool of four D50 COs was subsequently dissociated, and 2,427 single cells from the control and 1,655 from MCT8-deficient COs were processed for single-cell RNA-seq (scRNA-Seq; Parse) (Figure 1A).

**Figure 1.**
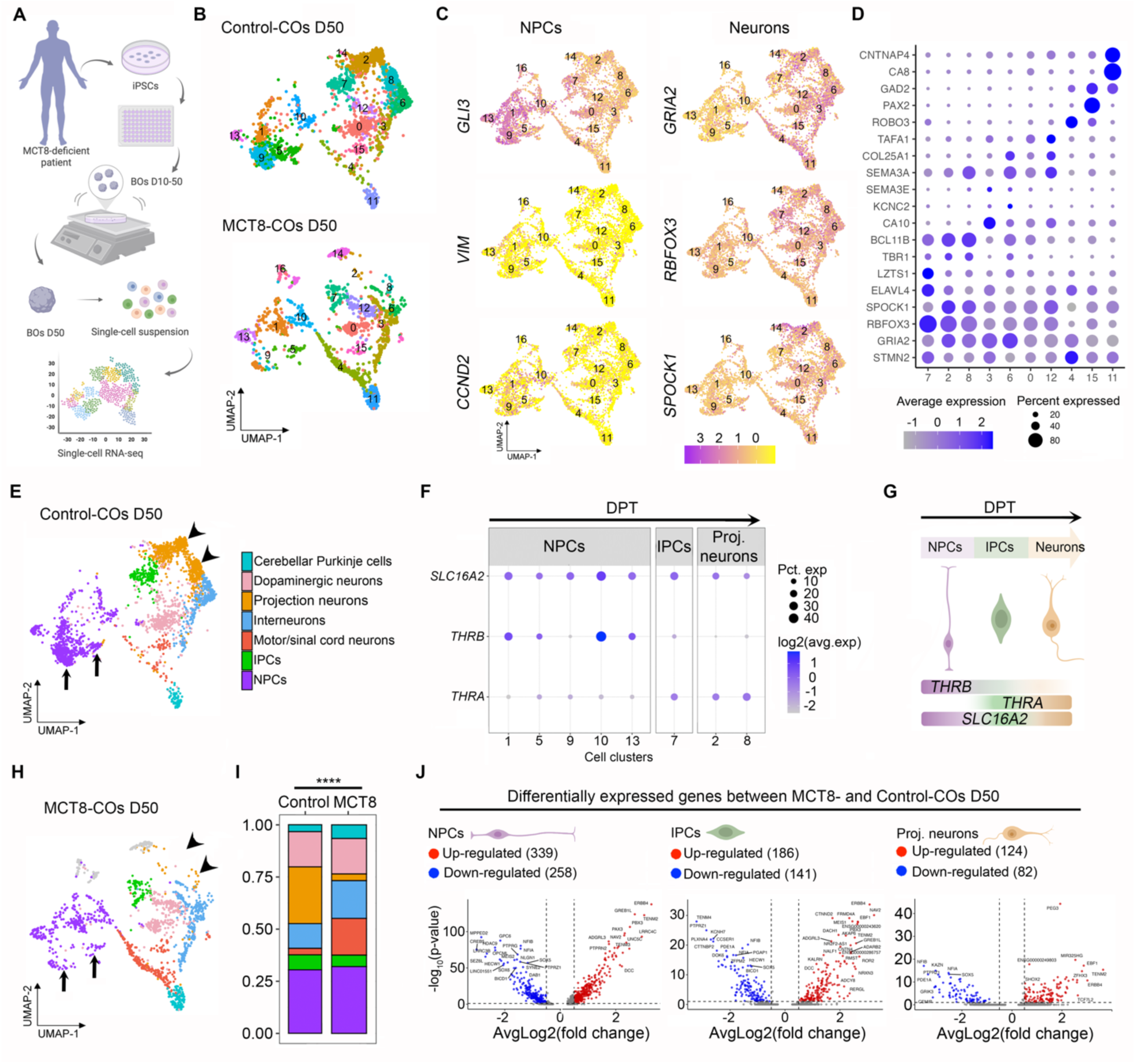
**A**. Schematic representation of generating D50 MCT8-COs and obtaining single-cell suspension for sc-RNA seq analysis. **B**. UMAP plot showing the cell clusters identified by principal component analysis in the indicated COs **C**. UMAP plots showing the distribution of markers for NPCs and neurons. **D**. Dot plot showing the relative expression levels of the gene markers used to identify each cell population. **E**. UMAP plot showing the cell types identified by manual curation in control COs. **F**. Dot plot showing the relative expression levels of SLC16A2, *THRB,* and *THRA* in each cell population of the dorsal projection trajectory. **G**. Interpretation of sequential gene expression changes during dorsal projection trajectory progression. **H**. UMAP plot showing the cell types identified by manual curation in MCT8-COs. **I**. Histogram of the relative number of cells in control and MCT8-COs. **J**. Volcano plots showing the distribution of differentially expressed genes in MCT8-deficient vs. control COs. iPSCs, induced pluripotent stem cells; COs, cortical organoids; NPCs, neural precursor cells; IPCs, intermediate precursor cells; DPT, dorsal projection trajectory; UMAP, Uniform Manifold Approximation and Projection. The X^2^ test for multiple comparisons and pairwise cell proportions. **P < 0.01; ***P < 0.0001. Differentially expressed genes thresholds: p-value < 0.05, and Average Log_2_ Fold-Change of 0.26.

Principal component analysis of the transcriptome and expectation-maximization clustering based on their position in the principal component space led to the identification of cell clusters that share genetic developmental markers (Supplemental Figure 1A-C; Supplemental Table 1), totaling 15 control and 16 MCT8-COs cell clusters (Figure 1B).

NPCs were contained in cell clusters 1, 5, 9, 10, and 13, expressing *GLI3*, *VIM*, and *CCND2* (Figure 1C). In turn, cluster 7 contained IPCs expressing *ELAVL4*, *LZTS1,* and low levels of *TBR1* (15–17), and the projection neurons were contained in clusters 2 and 8, expressing high levels of *TBR1, BCL11B, and GRIA2* (Figure 1C, D) (*18*). The other cell clusters contained neural cell types with non-dorsal projection trajectories, expressing the pan-neuronal markers *RBFOX3* and *SPOCK1* (Figure 1C) (19, 20). Among these non-dorsal neurons, clusters 3 and 6 contained interneurons expressing *CA10, KCNC2,* and *SEMA3E* (21), clusters 0 and 12 contained dopaminergic neurons expressing *SEMA3A COL25A1* and *TAFA1* (22, 23), clusters 4 and 15 contained motor/spinal neurons expressing *ROBO3*, PAX2, and *GAD2* (24, 25), and cluster 11 contained cerebellar Purkinje cells expressing *GAD2, CA8,* and *CNTNAP4* (26, 27) (Figure 1D, E).

To learn whether thyroid hormone signaling could influence the progression of the dorsal projection trajectory in our COs, we first looked for the expression of genes involved in thyroid hormone signaling and found that all three cell clusters belonging to the dorsal projection trajectory exhibited high *SLC16A2 expression* (encoding MCT8; Figure 1F). However, the expression of the thyroid hormone nuclear receptor isoform beta (*THRB*) and alpha (*THRA)* was heterogeneous amongst cell types. NPCs exhibited the highest expression levels of *THRB*, while IPCs and projection neurons mainly expressed *THRA*, altogether indicating that cells of the dorsal projection trajectory are equipped to respond to thyroid hormone (Figure 1G).

Indeed, the analysis of MCT8-COs versus control-COs revealed significant differences in the cellular composition of the cell clusters. The clusters of the dorsal projection trajectory were among the most affected (compare the UMAPs in Figure 1E with 1H). While MCT8-NPCs contained a similar relative number of cells, the subtypes of these cells among the different clusters were significantly different (X^2^ test, P < 0.0001, Figure 1I). As an example, NPC clusters 5 and 9 (arrows in Figure 1E, H) contained the largest number of cells in the control-COs but were markedly reduced in the MCT8-COs, revealing a very early deficit in the proliferation/differentiation capacity of NPCs (Supplementary Figure 1D). Furthermore, the IPCs and projection neurons in the MCT8-deficient COs had a reduction in the relative number of cells, with a ∼90% drop in the latter cells (X^2^ test, P < 0.0001; arrowheads in Figure 1E, H). In contrast, MCT8-COs exhibited an increased relative number of interneurons, motor/spinal neurons, and cerebellar Purkinje cell clusters with an altered distribution of these cells among clusters (X^2^ test, P < 0.0001; Supplementary Figure 1E). A corollary of these experiments is that the progression of the dorsal projection trajectory depends on an MCT8-mediated transport of thyroid hormone, impacting the final number of projection neurons in the developing fetal cerebral cortex.

To understand which aspects of the progression of the dorsal projection trajectory were affected by thyroid hormone, we next studied the differentially expressed genes between MCT8-CO and control-CO dorsal clusters and found 597 differentially expressed genes in NPCs (339 upregulated), 327 genes in IPCs (186 upregulated) and 206 genes in projection neurons (124 upregulated) (Figure 1J; Supplemental Table 2). The gene ontology analysis of MCT8-NPCs transcriptome identified 207 differentially expressed genes relevant to proliferation, regulation of neurogenesis, neuron differentiation, and axonogenesis. Furthermore, we also found differentially expressed genes involved in nuclear division, nuclear migration, and asymmetric cell division, which agrees with our previous observations that the plane of nuclear migration occurring during the translocation of the nuclei for mitosis is affected by a non-functional MCT8-transport of thyroid hormone (13). The differentially expressed genes found in the MCT8-IPCs belonged to 156 gene sets, including those regulating neuron projection development, neurogenesis, and neuron differentiation. Lastly, the differentially expressed genes in the projection neurons were contained in 82 gene sets, including those regulating glial cell differentiation, cell fate commitment, neuron differentiation and migration, and glutamatergic synapse (Supplemental Table 3). Overall, these studies provide an explanation for how an impaired thyroid hormone transport in MCT8-COs−which creates a state of localized hypothyroidism−leads to the arrested progression of the dorsal projection trajectory, resulting in a hindered differentiation of NPCs into projection neurons.

### The effects of thyroid hormone on NPCs, IPCs and neurons are independent of cell density

To obtain spatial information about NPCs, IPCs and neurons within the COs, we prepared two 5-um-thick slices from Control-COs and two from MCT8-COs D50 and analyzed them using a spatial molecular imager that provides transcriptomic data at a cellular resolution (Figure 2A, B). A universal cell characterization gene panel (Supplemental Table 4) was used to identify NPCs (expressing *VIM*, *NOTCH1*, and *BMP7*), IPCs (expressing *EOMES,* and *BMP5*), and neurons (expressing *GATA3*, *DCN*, *SPOCK2,* and *SOX4*) in control and MCT8-COs (Figure 2C, Supplementary Figure 2A). Undefined cells were considered altogether as a single group (Supplementary Figure 2B, C). The expression level of 72 genes was affected in the three groups of cells (Figure 2D; Supplementary Table 5). The top up- or down-regulated genes included *GPX1*, an antioxidant enzyme important for NPCs survival (28); *BCL2*, a protein that protects neural cells from apoptosis during early brain development (29); and *WNT7B,* a signaling protein that directs neuronal differentiation (30).

**Figure 2.**
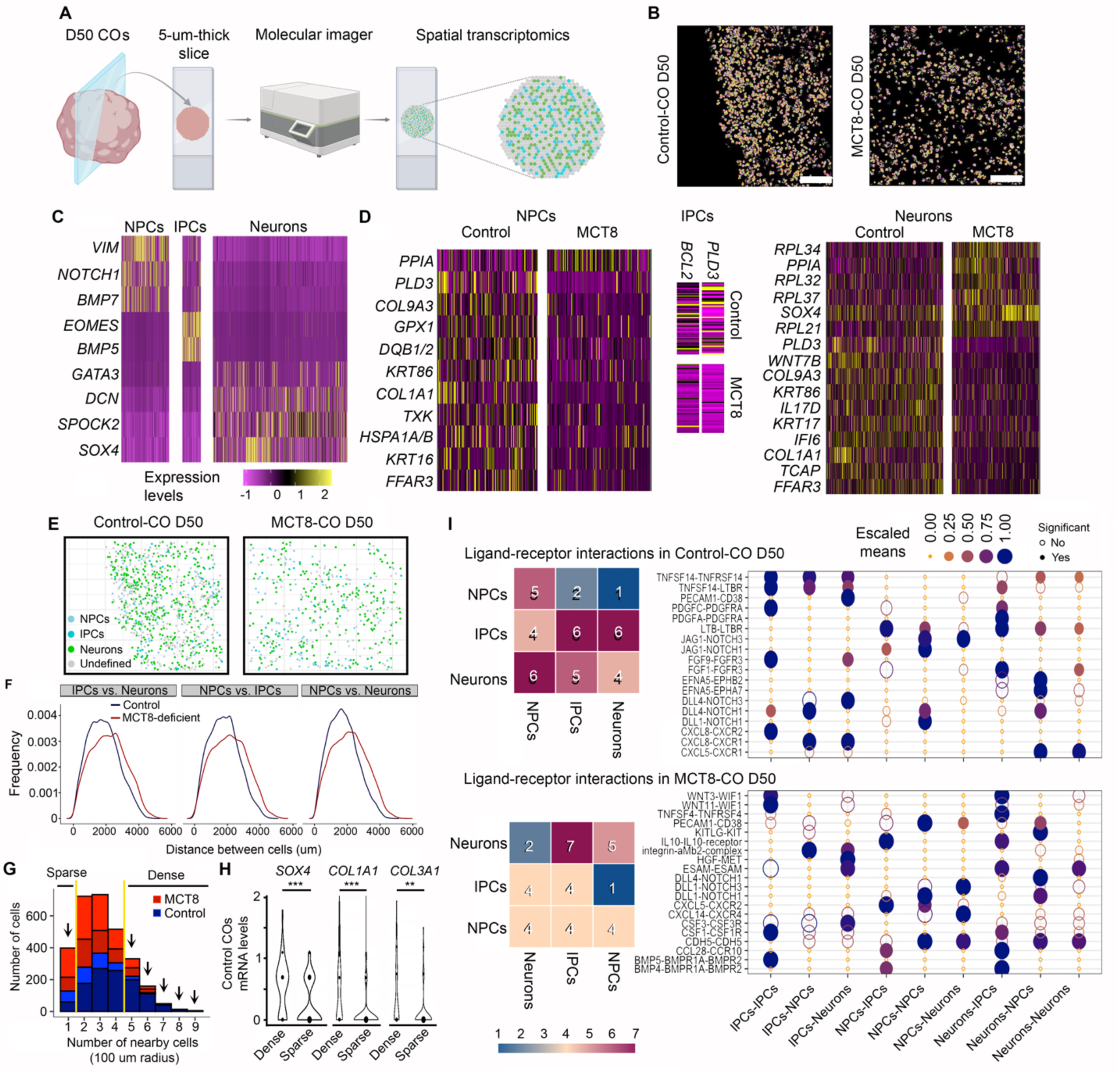
**A**. Schematic representation of obtaining spatial transcriptomic data. **B**. Image showing the cell segmentation on the indicated COs, based on transcript localization data. Scale bar: 100 µm. **C**. Heatmap showing the relative expression levels of the genes used to identify the indicated neural cell types. **D**. Heatmap of the differentially expressed genes in the indicated neural cells between control vs. MCT8-COs. **E**. Same as in **B**, except the cells are classified into the indicated types. **F**. Distribution of distances between the indicated cells and groups. **G**. Histogram of the distribution of cell densities. Considering a radius of 100 µm, if only one cell was in contact with another cell, it was considered as sparse density. If one cell was in contact with five or more cells, it was considered a dense density. **H**. Violin plots show the expression levels of the indicated genes, considering the indicated cellular densities. I. Ligand-receptor interaction between the indicated neural cells in Control and MCT8-deficient COs. Differentially expressed genes threshold: p-value < 0.05.

The analysis of the distance between cells revealed a higher density of cells (NPCs, IPCs, and neurons) in Control-versus the MCT8-COs (Figure 2E-F). To explore if the differences in cell density affected the transcriptome in Control- and MCT8-COs, we first identify two groups of cells that cluster together with sparse (cells with one cell in a 100 µm radius) and high densities (cells with five or more cells in a 100 µm radius), which resulted in 36 cells in sparse and 118 cells in dense clusters in the Control-COs and 31 and 71 cells in the MCT8-deficient COs, respectively (Figure 2G). Nonetheless, cell density only altered the expression of three genes (*SOX4*, *COL1A1* and *COL1A3*) in Control-COs (no genes were found altered in MCT8-COs) (Figure 2H).

Next, we used the CellPhoneDB, a bioinformatic toolkit designed to infer cell-cell communication (31, 32), to study the interactions of receptors and their respective ligands in nearby cells (receptor-ligand dyad). We found significant dyad differences between NPCs, IPCs and neurons in Control-COs and MCT8-COs (Figure 2I). In both COs, the most common dyad differences were the DLL1-NOTCH1 and DLL4-NOTCH1, which provide neural differentiation information to new neurons, and inhibitory signals that maintain the pool of NPCs (33). Only in Control-COs did we identify dyad differences between EFNA5-EPHB2 and -EPHA7, which are important for neuronal differentiation and axon guidance (34).

Overall, findings from the spatial transcriptomic analysis confirm the presence of NPCs, IPCs and neurons within COs. In addition, it confirmed major differences in the gene expression profile between Control and MCT8-COs. Furthermore, these differences were not related to cell density, reinforcing the effect of an impaired thyroid hormone transport on fetal neurogenesis.

### Thyroid hormone is critical for human NPCs to differentiate into neurons

To further explore the role of thyroid hormones on neurogenesis, we obtained a culture of iPSCs-derived NPCs (Figure 3A). Our iPSC-derived NPCs are immunoreactive for NESTIN, SOX2, and MCT8 (Figure 3B) and express the nuclear receptor isoforms *THRA* and *THRB* (Figure 3C), indicating they can respond to thyroid hormone. Therefore, we used media containing 20 pM free T3 to establish conditions under which 2D cultures of iPSC-derived NPCs generate cortical neurons in 12 days. The importance of thyroid hormone to the differentiation of NPCs became apparent with the observation that the ability of MCT8-NPCs to differentiate into neurons was impaired, even though they express *THRA* and *THRB*, albeit at lower levels. In MCT8-deficient NPCs, the expression of two markers of neuronal identity and maturity, *RBFOX3* and *NEUROD1*, remained steadily very low, indicating an arrested neurodifferentiation (Figure 3C).

**Figure 3.**
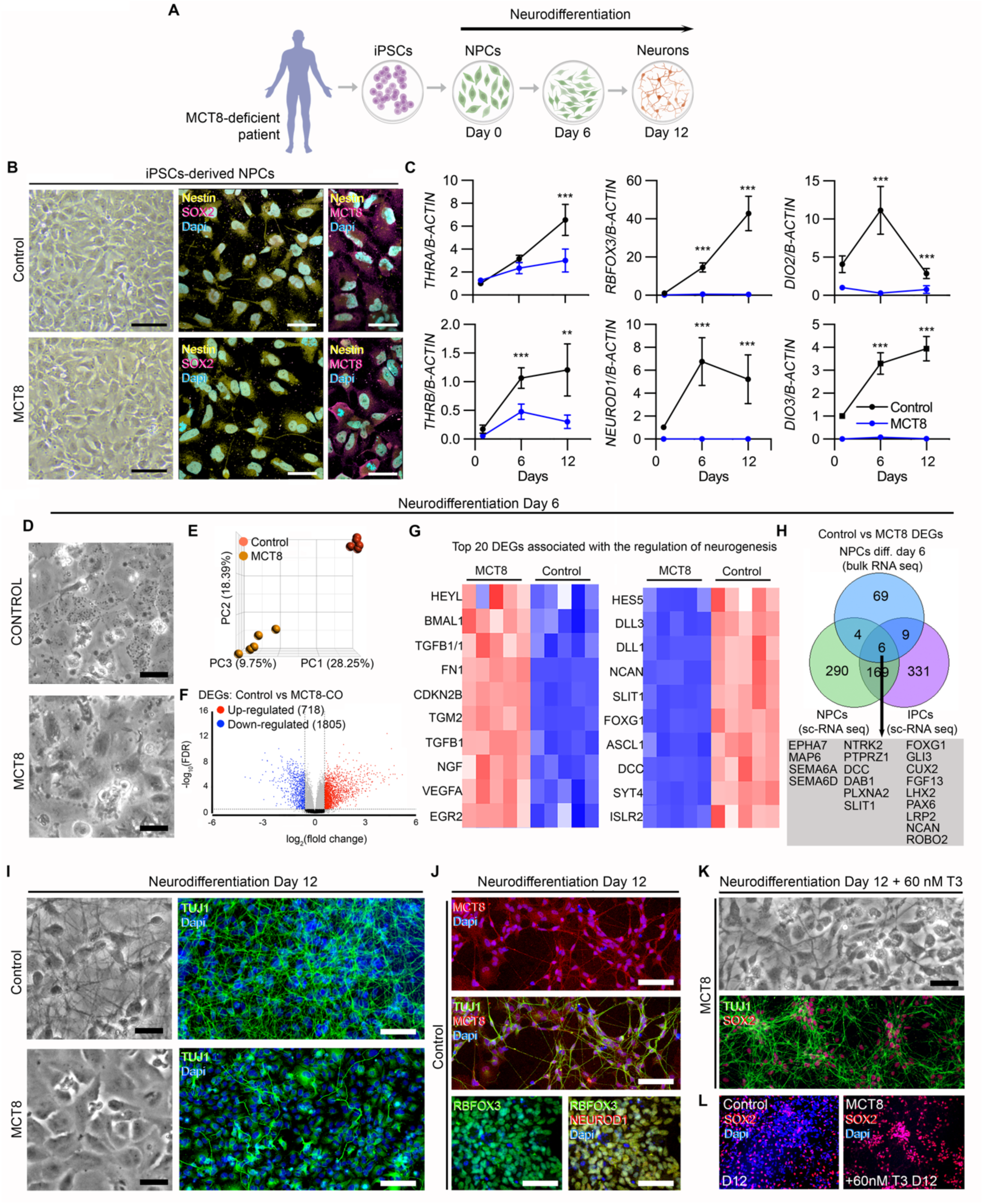
**A**. Schematic representation of generating NPCs from iPSCs and the subsequent differentiation of the NPCs into neurons. **B**. Brightfield and confocal fluorescence images showing iPSCs-derived NPCs stained for SOX2 (magenta), NESTIN (yellow), MCT8 (magenta), and Dapi (blue; nuclear). **C**. Relative mRNA levels of the indicated genes in Control and MCT8 NPCs after 1, 6, and 12 days of neurodifferentiation. **D**. Brightfield images showing NPCs after six days of neurodifferentiation. **E**. Principal component plot illustrating differences between control and MCT8-deficient NPCs. **F**. Volcano plots showing the distribution of differentially expressed genes in control vs. MCT8-NPCs; each point represents the average of 5 control and 5 MCT8-deficient samples of pooled NPCs for each transcript. **G**. Heatmap depicting the top 20 differentially expressed genes related to neurogenesis between control vs. MCT8-NPCs identified by bulk-RNA seq. **H**. Venn comparison of differentially expressed genes belonging to the gene set related to neurogenesis between control vs. MCT8-NPCs identified by bulk-RNA seq. (blue) and sc-RNA seq analysis (green) and between control vs. MCT8-IPCs identified by sc-RNA seq analysis (purple); common differentially expressed genes were identified (grey box). **I**. Brightfield images of control and MCT8-NPCs after twelve days of neurodifferentiation; TUJ1 staining in green and Dapi (blue; nuclear). **J**. The upper two panels are MCT8 staining in red, TUJ1 in green, and Dapi (blue); the lower panels are RBFOX3 staining in green, NEUROD1 in red, and Dapi in blue. **K**. MCT8-NPCs after being treated with 60nM T3 during the twelve days of neurodifferentiation. TUJ1 staining is green, and SOX2 is red. **L**. SOX2 staining in red and Dapi in blue on the indicated cells and treatments. Scale bars: B: 50 µm; D, I, J: 100 µm. Differentially expressed genes thresholds: p-value < 0.05, and Average Log_2_ Fold-Change of 1.5 in the Partek Flow platform. Expression values are mean ± SD of n = 3–6 RNA samples, each of them consisting of 2 pooled 6-well plates of NPCs from either control or MCT8-NPCs; Two-tailed Student’s test for comparing D2 deiodination and relative mRNA expression between D1, D6 and D12 of neurodifferentiation; *P < 0.05, **P < 0.01, ***P < 0.001.

Further analyses revealed additional mechanisms regulating thyroid hormone signaling during NPC differentiation. We detected a peak of the thyroid hormone-activating enzyme *DIO2* six days into the neuronal differentiation (6-day NPCs). This enzyme normally activates endogenous T4 to T3, transiently magnifying the T3 signaling (35). The peak of *DIO2* coincides with when the NPCs still exhibit normal morphology but are about to undergo major morphological changes typical of cortical neurons (Figure 3A, B, D). It is also notable that, all along the neuronal differentiation, there is a progressive increase in the expression of the thyroid hormone-inactivating enzyme *DIO3*. This is a typical neuronal marker that regulates access of T3 to the neuronal nucleus (36). In MCT8-NPCs, *DIO2* and *DIO3* levels also remained very low, indicating that changes in the expression of these enzymes are stage-specific.

We next process Control and MCT8 6-day NPCs for bulk RNA-seq analysis (these data sets clustered separately in a principal component plot; Figure 3D, E) and found major differences in their transcriptome, i.e. a total of 2,523 differentially expressed genes (1,805 downregulated in MCT8-cells) (Figure 3F; Supplemental Table 6). Through gene set analysis (Supplemental Table 7), we identified the top 10 up- or down-regulated genes associated with “regulation of neurogenesis” (Figure 3G), which include *HES5* (a notch effector during neuronal differentiation (37)) and *FOXG1*, a transcription factor critical for expanding the NPC pool (38). *FOXG1* heterozygous loss-of-function mutations cause FOXG1 syndrome, a severe neurological disorder where individuals frequently show absent speech, intractable seizure, and motor anomalies (39)–common alterations in patients with MCT8 deficiency (40). A top-upregulated gene in the MCT8-NPCs was *BMAL1*, a molecular clock that, when overexpressed, can lead to NPCs pool exhaustion (41). These findings indicate that the transcriptome changes triggered by impaired T3 signaling in MCT8-NPCs are substantial and associated with genetic programs driving neurogenesis and neuronal differentiation. Indeed, within the gene set “regulation of neurogenesis,” we identified 19 genes that were also differentially expressed in the NPCs and IPCs within MCT8-COs (Figure 3G green and purple circles, respectively).

After twelve days of culture, it became clear that a functional MCT8-mediated T3 transport is necessary to differentiate NPCs into neurons (Figure 3I). Indeed, only control cultures contained neurons that exhibited axons and other processes. Immunostaining for the postmitotic neuronal marker TUJ1 revealed robust neuronal processes in control neurons and short processes and projecting axons, all typically expressing MCT8 as described previously (Figure 3J; (36)). Neuron identity was further confirmed by colocalizing the neuronal markers RBFOX3 and NEUROD1 (Figure 3J). In contrast, neurodifferentiation was arrested in MCT8-NPCs, maintaining the NPC morphology with minimal TUJ1+ neuronal processes (Figure 3I).

These findings in human NPCs constitute a new model to test potential interventions that could rescue the devastating neurological phenotype seen in patients with MCT8 deficiency. With that in mind, we performed a proof-of-concept experiment in which the neural differentiation potential of the MCT8-NPCs was rescued by supplementing the medium with high concentrations of T3, which bypasses the MCT8 deficiency (60nM; Figure 3K). Remarkably, such treatment partially rescued the phenotype by promoting neuronal differentiation in MCT8-NPCs. Here, MCT8-neurons exhibited TUJ1+ neuronal processes (Figure 3K); however, all of these MCT8-neurons still retained a high expression of the NPCs maker SOX2 (vs. only ∼40% in control neurons; Figure 3L), suggesting that the MCT8-neurons, albeit partially rescued, were in a more immature state than controls. Overall, these findings indicate that the potential of NPCs to differentiate into neurons depends on an MCT8-mediated T3 transport and action.

### DIO2-generated T3 in human NPCs triggers genetic programs driving neurogenesis

DIO2 is active in D20 COs, a time when a high number of NPCs is expected (13). This led us to hypothesize that the resulting intracellular build-up of T3 could trigger developmental programs in these cells. To verify that NPCs exhibit DIO2 catalytic activity, we incubated 2D cultures of NPCs (obtained as in Figure 3A) for 24h with trace amounts of T4^I125^ (Figure 4A), which was deiodinated and led to a prominent peak of T3^I125^ (Figure 4B). As expected, T3^I125^ production was smaller in the MCT8-NPCs, indicating that T4 also uses MCT8 to enter NPCs (Figure 4B). Control-NPCs exhibit a DIO2 catalytic activity of 390 ± 140 pmol/mg T3/h, while MCT8-NPCs have a ∼4% of that in controls (37 ± 14 pmol/mg/h; Figure 4C).

**Figure 4.**
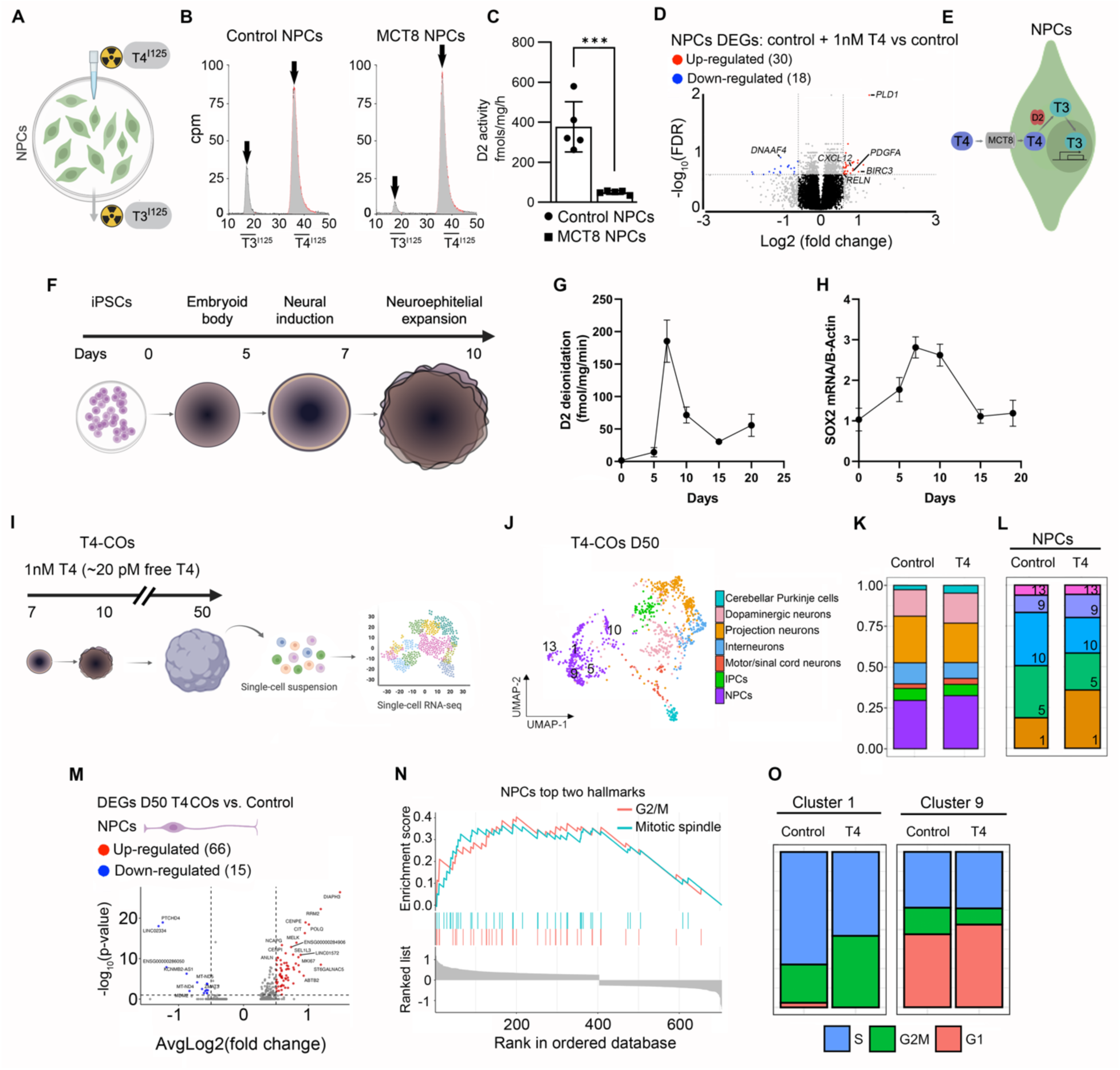
**A**. Schematic representation of treating iPSCs-derived NPCs with radioactive T4^I125^ to measure T3^I125^ production. **B**. Representative chromatograms of the medium after control and MCT8-NPCs were incubated with T4^I125^ for 24 hours. **C**. Quantitation of the DIO2 deiodination in control and MCT8 NPCs; n = 5 DIO2 assays. **D**. Volcano plots showing the distribution of differentially expressed genes in control + 1nM T4 vs. control NPCs; each point represents the average of five control + 1nM T4 and five control samples of pooled NPCs for each transcript. **E**. Interpretation of the findings in **A-D**. **F**. Schematic of the generation and timing of COs generation, starting with iPSCs to a culture of embryoid bodies, followed by neural induction, neuroepithelial bud expansion, and maturation. **G**. Quantitation of DIO2 deiodination in control COs during their first 20 days in culture. n = 4 DIO2 assays per timepoint, each consisting of 4 pooled COs from control COs. **H**. Relative *SOX2* mRNA levels in control COs during their first 20 days in culture. Expression values are mean ± SD of n = 3–6 RNA samples, each of them consisting of 4 pooled COs from control COs; **I**. Schematic representation of the experiment: COs are treated with 1nM T4 from D7 to D50 and then dissociated into a single-cell suspension for sc-RNA seq. **J**. UMAP plot showing the cell types identified. **K**. Histogram of the relative number of cells in T4-COs and control COs. **L**. Histograms of the relative number of cells in clusters of NPCs. The identification number of each cell cluster is indicated at the bottom right corner of each rectangle. **M**. Volcano plots showing the distribution of differentially expressed genes in T4-CO vs. control COs. **H**. Gene set enrichment analysis reveals gene ontology terms enriched in T4_TX_ COs. **O**. Histograms of the relative number of cells undergoing the indicated cell cycle phase in clusters one and nine of control and NPCs. Two-tailed Student’s test for comparing DIO2 deiodination in iPSC-derived NPCs; ***P < 0.001. Differentially expressed genes thresholds: p-value < 0.05, and Average Log_2_ Fold-Change of 0.26.

Based on other cell models (42), it is expected that substantial amounts of DIO2-generated T3 end up in the NPC nuclei, where it can then regulate (induction/repression) gene expression. To test if that was the case, control NPCs were incubated with 1 nM T4 (free T4 in the physiological range ∼20 pM; the DIO2 pathway is only relevant if T4 is supplied in the medium), and 24 hr later, NPCs were harvested and processed for bulk RNA-seq analysis (Supplementary Figure 4A).

Incubation with 1nM T4 resulted in 48 differentially expressed genes (30 upregulated) compared to the same cells not incubated with T4. (Figure 4D; Supplementary Table 8). The top gene *PLD1* encodes a phospholipase that is important in regulating the neuronal differentiation of NPCs (43). Another top gene, *BIRC3*, encodes an E3 protein ubiquitin ligase that impacts neural cell survival (44). The gene *PDGFA* was also found to be among the top upregulated genes by T4, acting as a mitogen for NPCs and stimulating their proliferation (45). The remaining genes included *DNAAF4*, which is associated with dyslexia and neuronal migration in the developing neocortex (46); *CXCL12*, controlling neurite outgrowth and axonal guidance and enhancing NPC cell survival (47); and *RELN*, a well-known T3-regulated gene that acts as a critical choreographer of neuronal positioning (48). The top gene sets with enrichment scores >6 were related to axon guidance, neuron migration, angiogenesis, cytoskeleton organization, forebrain cell migration, and neurogenesis (Supplementary Figure 4B).

Altogether, these results show that MCT8 function is critical for NPCs to take up T4 and— utilizing DIO2—activate T3 signaling (T4 + DIO2 → T3 + I^−^). The DIO2-generated T3 reaches the NPCs nuclei, where T3 triggers transcriptional changes related to cerebral cortex development, including neurogenesis (Figure 4E).

Next, we returned to the human cortical organoid model to study the role of the DIO2-generated T3 in NPCs in a more physiological context. First, we confirmed the presence of DIO2 catalytic activity during the initial 20 days of maturation, i.e. D0 (iPSC cells), D5 (embryonic bodies), D7 (neural induction), D10 (neuroepithelium expansion), D15 and D20 (maturation) (Figure 4F). We identified a peak of DIO2 catalytic activity at D7, reaching ∼180 fmol T3/h/mg protein (40-fold higher than D5), coinciding with the appearance of the first NPCs; DIO2 activity decreased afterward, stabilizing at values ∼10-fold higher than D5 (Figure 4G). Remarkably, a similar pattern was observed for the *SOX2* mRNA levels, a constitutive marker of NPCs (Figure 4H), suggesting that DIO2-T3 is driving the proliferation of the NPCs. The induction of high DIO2 activity at times when NPCs are highly proliferative (in a 3-month human embryo, NPCs proliferate at a rate of ∼ four million per hour (49)) suggests that the DIO2-generated T3 is invaluable for NPCs cell cycle.

To further explore this possibility, we studied developing COs that were treated with 20 pM free T3 + 20 pM free T4 (T4-COs) from D7 to D50 and compared the results with COs prepared in the presence of 20 pM free T3 (Figure 4I). After a pool of four D50 T4-CO was dissociated, 1,089 single cells were processed for scRNA-Seq, and cell clusters were identified as in Figure 1 (Figure 4J). The NPCs were the most affected by treatment with T4 (Supplementary Table 9). T4-NPCs clusters 1 and 9 exhibited a marked increase in the number of cells, while clusters 10 and 5 were significantly reduced, and 13 did not change (Figure 4L). These changes in the number of cells were paralleled by changes in gene expression so that T4-NPCs exhibited 81 differentially expressed genes (66 upregulated; Figure 4M).

Through gene set analysis, we identified that the top two main gene sets enriched in the T4-COs NPCs (same for clusters 1 and 9) were associated with cell cycle progression, regulating G2/M transition and the nuclear mitotic spindle (Figure 4N). One example of the T4-COs NPCs upregulated genes is *DIAPH3*, which encodes a protein that regulates the assembly and bipolarity of the mitotic spindle during neurogenesis (50). Another example is the gene *MELK*, which regulates the proliferation of NPCs (51). Further analysis confirmed that the treatment with T4 affected the NPCs cell cycle, increasing the number of cells in cluster 1 in the G2/M phase and the number of cells in cluster 9 in the G1 phase (Figure 4O). A corollary of these experiments is that physiological levels of T4 affect NPCs cell cycle and the plane of nuclear migration during the translocation of the nuclei for mitosis through the DIO2-generated T3.

### DIO2-generated T3 in mouse NPCs promotes neurodifferentiation

We next studied the effects of T4 and DIO2-generated T3 in a mouse model of differentiating NPCs. The advantage of this model is that the effects of T4 can be studied in the absence of T3 in the medium. The model was developed by establishing a protocol to isolate primary NPCs from embryonic day (E)13.5 mouse pups (mNPCs) (Figure 5A). mNPCs were isolated as neurospheres in serum-free medium that contained the in-house made supplement B26 (similar to B27 except that it does not contain T3 (52)) in the presence of epithelial growth factor (EGF) (Figure 4B). After dissociation, mNPCs exhibited their characteristic apical and basal cellular processes and expressed nestin and sox2 (Figure 4C, D) as well as Mct8, exhibiting a more intense immunofluorescence signal in the nuclei of mNPCs undergoing cell divisions (Figure 4E). mNPCs exhibited DIO2 catalytic activity (∼6.5 fmol T3/h/mg protein) (Figure 4F). These findings indicate that mNPCs are equipped with the MCT8-DIO2 dyad, which allows them to take T4 and build up intracellular levels of T3 to enhance TH signaling.

**Figure 5.**
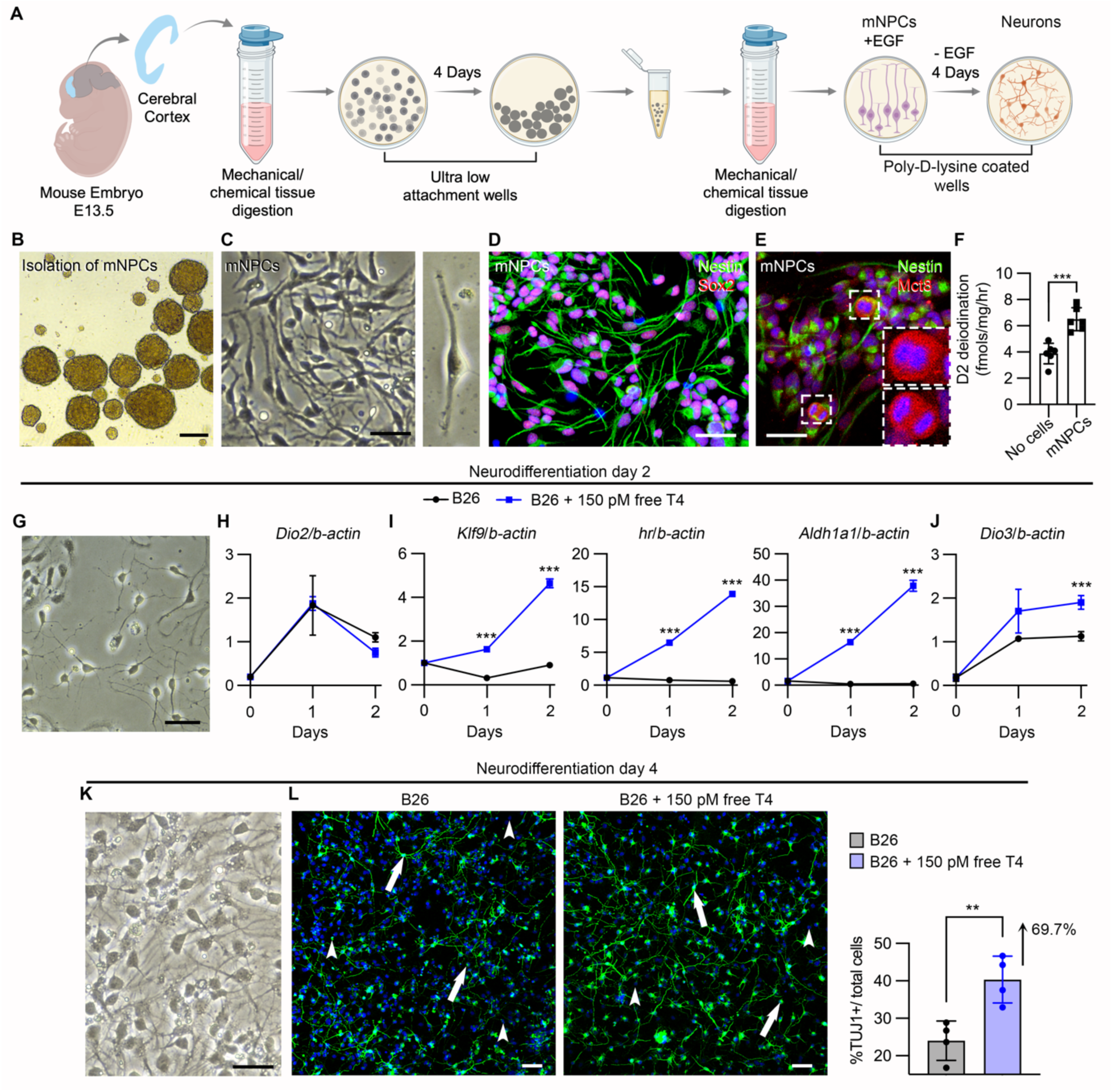
**A**. Schematic representation of the protocol to isolate mNPCs, propagate them in culture, and differentiate into neurons. **B, C.** Brightfield images showing representative neurospheres containing mNPCs (**B**) and the mNPCs cultured in collagen-coated plasticware (**C**). **D-F**. Confocal images showing mNPCs expressing Nestin (green) and Sox2 (red) (**D**), and Mct8 (red) (**E**). The insets in **E** depict two dividing mNPCs that exhibited higher intensity of Mct8 immunofluorescence. **F**. Quantitation of the DIO2 deiodination in mNPCs; n = 6 DIO2 assays. **G.** Brightfield images showing representative mNPCs after two days of neurodifferentiation. **H**-**J**. Relative mRNA levels of the indicated genes in mNPCs under the indicated conditions; n = 4. **K**. Brightfield images showing representative mNPCs after four days of neurodifferentiation. Note the neuronal process extension. **L**. Tuj1+ cells under the indicated conditions and the quantitation of the percentage of Tuj1+ cells. Values are the mean ± SD of 5 replicates. Scale bars: B: 300 µm, C-L: 25 µm. Two-tailed Student’s test for comparing DIO2 deiodination in iPSC-derived NPCs; **P < 0.01; ***P < 0.001.

The differentiation of mNPCs into neurons was started by removing EGF from the medium, and as soon as after 48 hours, mNPCs transitioned to a more neuron-like morphology with small cellular processes (Figure 5G). We documented that during this initial 48 h of neurodifferentiation, mNPCs exhibited an increase in the *Dio2* levels, peaking at 24 h and decreasing afterward (Figure 5H), indicating that during this period of neurodifferentiation, mNPCs can activate T4 into T3 via the Dio2 pathway. Therefore, we used media containing ∼150 pM free T4 to establish conditions under which mNPCs can respond to T4 while undergoing neurodifferentiation. mNPCs treated with T4 exhibited similar levels of Dio2 (Figure 5H), but the expression of *Klf9*, *Hairless*, and *Aldh1a1,* three known T3-regulated genes (53), markedly increased in the mNPCs treated with T4 (Figure 5I). As expected, during the 48 h of neurodifferentiation, there was a progressive increase in the expression of the TH-inactivating enzyme Dio3 (Figure 5J), which is normally expressed in neurons. Considering the important role played by *Klf9*, *Hairless*, and *Aldh1a1* during neurodifferentiation (54, 55), these findings confirm a role for D2-generated T3 in this process. Indeed, after 96 h of neurodifferentiation (Figure 5K), the number of tuj1-positive neurons in the cells treated with T4 was 67.9% higher (Figure 5L). A corollary of these experiments is that the DIO2 pathway (T4 + D2 → T3 + I−) in mouse NPCs promotes neurodifferentiation.

## DISCUSSION

Our results obtained in human cortical organoids and in primary cultures of mouse NPCs uncover several key mechanisms by which thyroid hormone regulates neurogenesis during the fetal cerebral cortex development. First, thyroid hormone is important in the progression of the dorsal projection trajectory during the first trimester of the fetal cerebral cortex development. iPSC-derived MCT8-NPCs not only presented dramatic changes in the expression of critical genes for regulating fetal neurogenesis and neuron differentiation but also failed to differentiate into neurons. Publicly available datasets from human and murine cells allocate *DIO2* to NPCs (11, 19, 56). Here we used iPSC-derived NPCs and further revealed that the DIO2 pathway is catalytically active and can enhance TH signaling by activating T4 to T3 in these cells. This pathway triggers transcriptomic features that are pro-neurogenesis and important for NPC cell cycle progression, effectively promoting neuronal differentiation. Our results not only identified the role of the MCT8-DIO2 dyad in the development of the dorsal projection trajectory but also provide insights into the mechanisms that allow T4 to act during the early stages of fetal cerebral cortex.

The appearance of thyroid hormone receptors (TRs), alpha (TRα), and beta (TRβ) during the development of the fetal cerebral cortex marks the moment that thyroid hormone signaling becomes essential in this process. TRs appear during gestational weeks 7-8, increasing their levels (protein and mRNA) tenfold from weeks 10 to 16 (57). It is well-accepted that TRα is important because *THRA* mutations are more frequently associated with a neurological phenotype than *THRB* mutations. But about one-half of patients with *THRB* mutations have learning disabilities, and a low IQ (<60) can be present in ∼3% of cases (58); ∼50% of the children carriers of TRβ mutations are diagnosed with attention deficit hyperactivity disorder (59).

Our COs modeled the first trimester of human fetal brain development (6.5 to 14 gestational weeks) (60), revealing a shift in the expression of the predominant TR in the dorsal projection trajectory. First, *THRB* is expressed in NPCs, followed by *THRA* in IPCs and projecting neurons. In agreement, affinity studies show that the fetal TRs exhibit a higher affinity for the TRβ-selective agonist TRIAC than for T3 (57), suggesting that most fetal brain TRs at gestational week 10 to 16 are TRβ (61). In addition, a compilation of studies indicates that T*HRB* is important for the development, migration, and function of interneurons (11, 62, 63). Other studies in COs from iPSCs of humans and gorillas have shown an increase in the expression of *THRA* (isoform 2) during the dorsal projection trajectory, being low in NPCs and progressively increasing to a peak in neurons (64). Unfortunately, the expression of *THRB* was not reported in these studies.

Our previous studies using MCT8-deficient COs and primary cultures demonstrated that MCT8 mediates the bulk of T3 transport in developing neural cells (13) and neurons (36), providing evidence that the role of this transporter in the human brain goes well beyond facilitating the passage of TH through the blood-brain barrier. The present investigation provided yet an even clearer understanding of the role of MCT8-mediated transport of TH on cerebral cortex development.

First, our results support the hypothesis that thyroid hormone acts as a molecular cue that helps define the fate of specific neuronal cell types. This is illustrated by the selective significant reduction in the number of projection neurons observed in MCT8-COs, while other neuronal types remained unaffected. It is then conceivable that part of the severe neurological phenotype present in patients with MCT8-deficiency can result from a severe imbalance in the distribution of neuronal cell types in different brain regions without altering the total number of neurons. In agreement, a compilation of studies in animal models supports the idea that an optimal MCT8-mediated transport of thyroid hormones influences the balance between inhibitory and excitatory neurons (62, 63, 65, 66). Further evidence comes from a study that measured the incorporation of the tracer ^13^C-glucose in excitatory and inhibitory neurotransmitters in adult hypothyroid mice and found a higher incorporation of the tracer in the inhibitory neurotransmitters (67). Future studies using Magnetic Resonance Spectroscopy could measure the excitatory neurotransmitter glutamate and the inhibitory gamma-aminobutyric acid in the brains of patients with MCT8 deficiency to help clarify whether there is an imbalance between excitatory and inhibitory neural activities in the brains of these patients.

Second, we found that thyroid hormone signaling contributes to the ability of NPCs to differentiate into neurons. This is illustrated in the experiment in which we discovered that MCT8-NPCs exhibit down-regulated gene sets related to neurogenesis and neuronal differentiation and by the fact that MCT8-NPCs failed to differentiate into neurons. These results expand on our previous observation that genes involved in neurogenesis and neuronal differentiation are reduced in D65 MCT8-deficient COs (13) and are further supported by a study analyzing brain and cerebellum sections from an MCT8-deficient fetus and identifying abnormalities in the density of neurons (68). Similar evidence was found in MCT8-knockin mice (P235L), which exhibit fewer neurons in layers I–IV of the somatosensory cortex (projection neurons are the predominant neurons in layers II-IV) (66).

The present investigation gave us a much better insight into the mechanistic underpinnings of how T4 works during early cerebral cortex development. It was known that the human NPCs expressed *DIO2* (11), and we previously described DIO2 activity in COs in a period when NPCs are very active (13). Here, we complete the picture and show that NPCs indeed can take up T4 and activate to T3 via the DIO2 pathway. In addition, we also demonstrated that MCT8 is responsible for the bulk of the T4 transport across the membranes of the NPCs. This is illustrated by the reduced DIO2-mediated T4^I125^ to T3^I125^ conversion in MCT8-NPCs. The peak of T3^I125^ indicates the uptake of T4^I125^ into the NPCs, its metabolism, and the release of T3^I125^ to the medium.

The T3 generated by DIO2 can alter the transcriptome of NPCs cultured in 2D and within COs. It is important to note that many of the genes regulated by T4 (via the DIO2 pathway) are involved in critical processes of NPCs biology, and the alterations of only a few of them can result in severe consequences; the control of *DIAPH3* (top up-regulated) is a good example. *DIAPH3* is exclusively expressed in NPCs and is a major regulator of the actin cytoskeleton (69). The lack of DIAPH3 compromises nuclear division in NPCs, resulting in mitotic errors, defective neurogenesis, and, ultimately, brain dysfunction (50). In agreement, studies on rats show how pups exposed to low thyroid hormone levels during pregnancy present impaired maturation of NPCs (70). Considering that (i) DIO2 acts as a dynamic switch (71) to trigger developmental transitions in the cerebellum (10), retina (72, 73), cochlea (74), brown adipose tissue (75), and liver (76, 77) and that (ii) DIO2-generated T3 seems to be important for the cell cycle of NPCs, our study supports the idea that DIO2 could also act in NPCs as a switch to control the cell cycle progression of these cells. Indeed, T4 treatment changed the proportion of NPCs on different cell cycle phases: some NPCs exhibited shorter G2/M phases and others longer; such changes in G2/M phases are associated with more proliferation or neurodifferentiation, respectively (78).

The limitations of the study include the fact that we assessed the effect of thyroid hormones on COs only through transcriptional profiles. Furthermore, because no datasets of human fetal cortex cover the entire span of cortical development, it was difficult to establish a time relationship between the D50 COs and the human fetal cortical development. Finally, we utilized only two iPSC lines, the control line was obtained from the unaffected father of the patient with approximately 50% genetic similarity, which has likely minimized the genetic background variance. Future studies should consider using the currently available array of iPSCs derived from patients with thyroid genetic conditions, including mutant *THRB*, *THRA*, and *SLC16A2* (encoding MCT8) (79–82), to generate cortical organoids and explore mechanisms of thyroid hormone action in fetal cerebral cortex development.

In conclusion, we have identified intracellular mechanisms in human NPCs that can transport T4 and activate it into T3. The mechanisms involve, respectively, two components: the TH transmembrane transporter MCT8 and the enzyme DIO2. These components work together to rapidly enhance TH signaling by taking and building up locally generated T3 in the NPCs. The MCT8-DIO2 dyad customizes T4 signaling and regulates NPCs’ proliferation and neurodifferentiation potential. Remarkably, the progression of the dorsal projection trajectory (i.e., NPCs → intermediate progenitors → projection excitatory neurons) is dramatically affected by a suboptimal transport of TH. In other words, this work constitutes objective evidence that at least one trajectory of neurons during cerebral cortex development may be compromised in low T4 conditions. This new observation warrants reassessing the relevance of low T4 levels for cerebral cortex development, which often affects normal pregnancies and preterm infants. In addition, this research contributes to our understanding of the pathophysiology of MCT8-deficiency, a condition that significantly hinders neurodevelopment and for which treatment continues to be understandably difficult.

## Methods

### Sex as a biological variable

Our study exclusively examined cell lines derived from male patients and one from his father because the disease modeled is only relevant in males.

### Cell lines and maintenance

iPSC lines were obtained and maintained as previously described (13). We used a cell line from an MCT8-deficient subject (CS58iMCT8) and one line derived from his father (CS01iCTR) to serve as a control. The karyotype and pluripotency of each iPSC line were verified as previously described (13, 82).

### Generation of human cortical organoids

COs were generated and cultured as previously described (13). Briefly, on D0, 80% confluent human iPSCs were dissociated using the Gentle Cell Dissociation Reagent, and approximately 9,000 cells/well were seeded in a 96-well, ultralow-attachment plate (Corning) in EB formation medium containing 10 μM Y-27632 (Tocris). On D5, the medium was removed, and EBs were incubated with a neural induction medium. On D7, EBs were kept on the 96-well plate, and the medium was supplemented with 2% Matrigel. On D10, EBs were transferred into ultra-low-attachment, 6-well plates kept in a maturation medium on an orbital shaker (75 rpm), with media changes every other day. From D20 onward, the maturation medium was switched to COs maturation medium containing BrainPhys and the supplements SM1, N2A, NEAA (Gibco), Glutamax (Gibco), insulin (MilliporeSigma), BME (MilliporeSigma), and 2 ng/mL brain-derived neurotrophic growth factor (Tocris). Every 3 days, half of the medium was replaced with a fresh one.

### Single-cell RNA sequencing

The libraries were prepared as in (52, 77), using the Parse Bioscience kit and were sequenced using a HiSeq Illumina platform, paired-end setting. The spipe pipeline from Parse Biosciences was run in mode all and combined for eight samples, with a total of 10719 cells. Further processing was done in R using Seurat v5. Initial QC was performed by sample, filtering for cells with 500 to 10,000 genes detected; total transcript counts between 1,000 and 15,000; less than 10% of mitochondrial transcripts and removing for doublets after using scdblfinder, accounting for 8055 cells. We performed integration using Seurat using the top 2000 most variable genes per sample. After integration we performed principal component analysis, using the first 15 principal components for UMAP dimension reduction. We performed SSN clustering at 0.8 resolution. Cluster annotation was performed using sc-type, using as reference the Human, brain dataset. After selecting the cell lineages of interest, we redid non-linear dimensional reduction by Uniform Manifold Approximation and Projection and clustering by the shared nearest in the subsets. The differential expression analysis was computed using the DeSeq2 and MAST methods in the Seurat environment. We used a p-value < 0.01 and fold change > 1.5 as thresholds for differentially expressed genes, comparing MCT8-COs against Control-COs and Control-COs against T4-COs. Gene set enrichment analysis was performed using clusterProfiler.

### Spatial Transcriptomics

After producing Flatfiles (Check the methods from the other lab to get these files), we process the data using Seurat. Initial QC was performed with a relax filtering, filtering for cells with more than 15 probes with transcripts and more than 20 transcripts. This reduced the set from 3284 cells to 2928 in the final set. Data was normalized using all the 1207 genes in the probe set. PCA was calculated, and the first 10 PCs were used for subsequent UMAP and clustering, with an 0.8 SNN resolution. Cell annotation was performed using a set of gene markers, defining cells that expressed at least one of the gene markers for a cell type and none of the other markers as part of the cell type set. For the analysis of the distance between cells, we calculated the Euclidean distance between cells of different cell types per sample. For a given pair of cell types, we aggregated the distances by genotype afterward. To test whether there was a difference in the distributions of distances of cells from different cell types between genotypes, we performed the Kolmogorov-Smirnov test to compare the distributions. The cell density, or cell neighborhood metric (n) defined as the number of cells within a 100 µm radius from a given cell. We tested the effect of cell density in gene expression per cell type using linear regression, using n metric as the covariate, and logistic regression using as categorical variable whether a cell was in sparse areas (Cells with n==1) or dense areas (Cells with n>=5). We performed multiple testing p-value adjustments using the Benjamini-Hochberg method. The analysis of the differentially expressed genes was performed using the findMarkers function and Wilcox test as method for testing within each cluster for the Parse dataset or cell type group for the CosMx dataset, comparing between genotypes or treatments. Differentially expressed genes were selected as those with an adjusted p-value under 0.05, and for Average Log_2_ Fold-Change, no threshold for CosMx data and a threshold of 0.26 for the Parse dataset. Gene set enrichment analysis for gene ontology terms, and gene set enrichment analysis were performed using clusterProfiler in R.

### Generation and culture of NPCs from iPSCs

On day 0, 80%-confluent iPSC were dissociated using the Gentle Cell Dissociation Reagent (STC 100-1077), and 2 x 10^6^ cells/well were seeded in a single well of a matrigel-coated 6-well plate (Corning) in 2ml of Neural Induction Medium (SCT 08581) containing inhibitors of the SMAD signaling and 10 µM Y-27632 (Tochris). From this point on, the medium was changed daily (without Y-27632), and on day 7, cells were detached using ACCUTASE and passed to a new matrigel-coated 6-well plate (1 to 3 ratio). After three passages, cells were cultured in Neural Progenitor Medium (SCT 05833; the medium was replenished daily) for 1 – 3 days until cells were 50% confluent and ready for the neuronal differentiation protocol.

### Neuronal differentiation of iPSCs-derived NPCs

When NPCs were at 50% confluency, we added an equal volume of differentiation medium composed of BrainPhys (SCT 05790), 2.5%, SM1 (SCT 05711), 1% N2-A (SCT 07152), BDNF (20ng/ml; Tochris), GDNF (20ng/ml; Tochris), and L-ascorbic acid (200nM; Tochris). Every 2 - 3 days, ∼50% of the differentiation medium was removed from each well and replaced with an equal volume of fresh medium. A 10 to 15 days culture period was required for cells to differentiate and exhibit neuronal morphology.

### Immunofluorescence studies

iPSC-derived NPCs and neurons were fixed in 4% paraformaldehyde for 20 min, washed twice in PBS, and then permeabilized in PBS with 0.1 % Triton X-100 for 5 min. Fixed cells were then incubated with primary antibody diluted in PBS, at 4°C overnight, rinsed 3 times with PBS for 20 minutes and incubated for 2 h at room temperature with a secondary antibody. Mouse monoclonal anti-Nestin antibody 1:1000 (Biotechne; AF4320), rabbit polyclonal anti-MCT8 antibody 1:400 (Atlas antibodies; HPA003353), mouse monoclonal anti-Tuj1 1:1,000 (Bio-Techne; AF4320), anti-Sox2 1:1,000 (Abcam; 109186), guinea pig polyclonal anti-Rbfox3 (Millipore; ABN90). Alexa Fluor 488– conjugated goat anti-mouse IgG 1:200 (Vector Laboratories; DK-2488), Alexa Fluor 594– conjugated horse anti-rabbit IgG 1:200 (Vector Laboratories; DK-1594), and Alexa Fluor 488– conjugated horse anti-rabbit IgG 1:200 (Vector Laboratories; DI-1788). Nuclei were counterstained with DAPI (1:10,000; Invitrogen; D1306). Alexa Fluor 488–conjugated horse anti-guinea pig IgG 1:200. The images were analyzed with a NIS-Element AR (Nikon Instruments) and final figures were prepared for publication on Adobe Photoshop.

### Iodothyronine chromatography using ultra-high-performance liquid chromatography

NPCs were processed for DIO2 assays as previously described (83, 84). Before starting the DIO2 assays, T4^I125^ was purified on an LH-20 column. NPCs were incubated with 250,000 cpm of T4^I125^/ml in the presence of 10 nM T3 to saturate DIO3. After 24, 100 ul of the medium was sampled, mixed with 100 ul of 0.02 M ammonium acetate + 4% methanol + 4% PE buffer (0.1M PBS, 1mM EDTA), and applied to the UPLC column (AcQuity UPLC System, Waters). Fractions were automatically processed through a Flow Scintillation Analyzer Radiomatic 610TR (PerkinElmer) for radiometry. The DIO2-mediated deiodination was normalized to the protein concentration (bicinchoninic acid method; Pierce, Thermo Fisher Scientific) and expressed as fractional conversion (fmol/mg/h or fmol/mg/min).

### TaqMan Real-Time quantitative PCR

Total RNA was extracted from ∼1X10^6^ neural cells per group and stage of differentiation. mRNAs were treated with DNase (QIAGEN) and measured by quantitative real-time PCR (85). Briefly, total RNA was isolated using a QIAGEN RNeasy Mini Kit, according to the manufacturer’s instructions. The cDNA was prepared using a cDNA synthesis kit (Roche). Data were analyzed using the 2^−ΔΔCT^ method and displayed relative to an arbitrary value. The expression of the indicated genes was determined using specific primers. The expression of ß-ACTIN was used as the internal control.

### RNA sequencing and analysis

Samples of total RNA were sent to the Genomic facility at the University of Chicago for library preparation and sequencing. Libraries were paired-end sequenced with NovaSeq S4 (Illumina). Base calls and demultiplexing were performed with Illumina’s bcl2fastq software and a custom python demultiplexing program with a maximum of 1 mismatch in the indexing read. The FASTQ files were aligned to gencode hg38 transcriptome with STAR (v.2.7.8a) using the Partek Flow platform. Aligned reads were quantified to the annotation model (Partek E/M) and normalized to counts per million.

### Isolation of NPCs from mouse embryonic cerebral cortices and differentiation into neurons

mNPCs were isolated from E13.5 mouse embryos. Briefly, embryos were removed, and the cerebral cortex dissected, stripped of meninges, and dissociated into a combination of Ca2+ and Mg2+ free Hanks balanced salt solution (HBSS) containing ACCUTASE, then mechanically triturated using fire-polished glass Pasteur pipettes. Isolated cells were passed through a 40 μM cell strainer and were pellet by centrifugation (1500 rpm/5min). Cells were resuspended in mNPCs complete medium composed of DMEM/F12, 2% B27, 1% antibiotic-antimycotic (Penicilin-streptomycin; all from Gibco), and epithelial growth factor (EGF; 20ug/ul; R&D Systems 2028-EG-200). For experiments in which the mNPCs were incubated with T4, B27 was replaced by B26, an in-house supplement that does not contain T3 (52, 77).

### Statistics

All data were analyzed using Prism software (GraphPad). Unless otherwise indicated, data are presented as scatterplots depicting the mean ± SD. Comparisons were performed by a 2-tailed Student’s t test and multiple comparisons by 1-way ANOVA followed by Tukey test. A P < 0.05 was used to reject the null hypothesis.

### Study approval

All experiments were approved by the Institutional Animal Care and Use Committee at the University of Chicago (#72577) and followed the American Thyroid Association Guide to investigate the TH economy and action in rodents and cell models (86).

## Authors contributions

FSL conceptualized the study, conducted experiments, prepared figures, analysed and interpreted the data, and prepared the manuscript. SE conducted experiments and prepared supplemental materials. HJ prepared the figure schematics, AC and RS obtained the molecular imager data. ACB interpreted the data and edited the manuscript. FSL directed all the studies.

## Acknowledgments

The authors thank Diana Alvarez, PhD, for her help analyzing the transcriptomic data. Authors are grateful for support from NIH National Institute of Diabetes and Digestive and Kidney Diseases DK15070 and DK65055. F.S.-L. was partly supported by the American Thyroid Association research grant #4634920.

## Notes

### Competing Interest Statement

A.C.B. is a consultant for AbbVie, Acella, Aligos, and Synthonics. The other authors have nothing to disclose.

