## Supplemental Figures for "Thyroid hormone promotes fetal neurogenesis"

### Supplemental figure legends

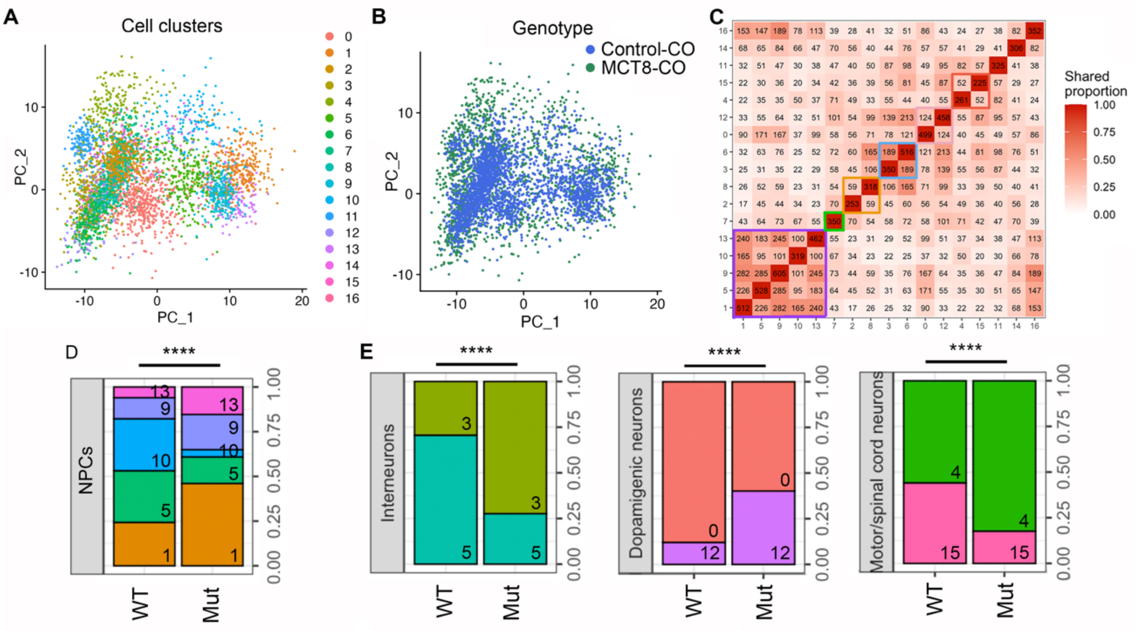


**Supplemental Figure 1. A**. PCA of the two first principal components (PC1 versus PC2) of D50 COs. **B**. Same as in **A**, except that the different genotypes are indicated. **C**. A plot of the number of conserved genes shared across the clusters. Coloured squares (purple, green, orange, blue, and red) indicate the cluster of cells for cells belonging to different cell types. **D**. Histograms of the relative number of cells in clusters of NPCs. The identification number of each cell cluster is indicated at the bottom right corner of each rectangle. **E**. Same as in **D** for the indicated type of neurons. The $\chi^2$ test for multiple comparisons and pairwise cell proportions. ***P < 0.0001.

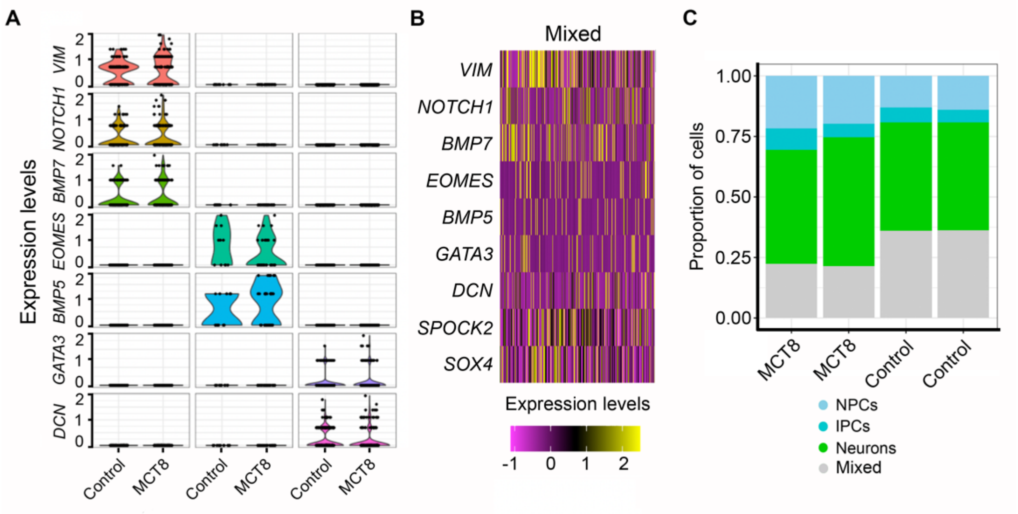


**Supplemental Figure 2. A**. scRNA-Seq violin plots of the expression levels of the indicated markers and the indicated groups of COs. **B**. Heatmap showing the relative expression levels of the cell identity marker genes in the group of mixed cells. **C**. Histograms of the relative number of cells in the indicated cells and groups of COs.

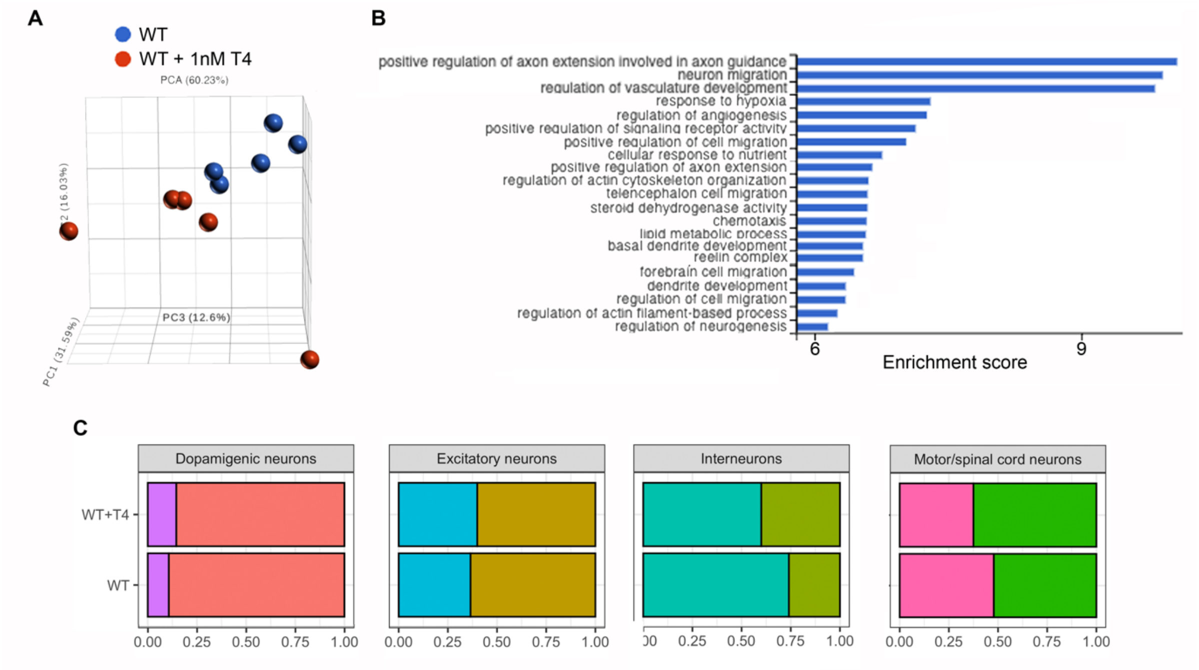


**Supplemental Figure 3. A.** Principal component plot illustrating differences between control and control + 1 nM T4 NPCs. **B**. Top gene sets with enrichment scores >6. **C**. Histograms of the relative number of cells in the indicated type of neurons.
